# Insight into mammary gland development and tumor prevention in a newly developed metastatic mouse model of breast cancer

**DOI:** 10.1101/2021.09.24.461727

**Authors:** Briana To, Carson Broeker, Jing-Ru Jhan, Rachel Rempel, Jonathan P. Rennhack, Daniel Hollern, Lauren Jackson, David Judah, Matt Swiatnicki, Evan Bylett, Rachel Kubiak, Jordan Honeysett, Shams Reaz, Joseph Nevins, Eran Andrechek

## Abstract

The development of breast cancer has been observed due to altered regulation of mammary gland developmental processes. Thus, a better understand of the normal mammary gland development can reveal possible mechanism in how normal cells are re-programmed to become malignant cells. E2F1-4 are part of the E2F transcription factor family with varied roles in mammary development. However, little is known about the role of E2F5 in mammary gland development. A combination of scRNAseq and predictive signature tools demonstrate the presence of E2F5 in the mammary gland and showed altered activity during the various phases of mammary gland development and function. Testing the hypothesis that E2F5 regulates mammary function, we generated a mammary-specific E2F5 knockout mouse model, resulting in modest mammary gland development changes. However, after a prolonged latency the E2F5 conditional knockout mice developed highly metastatic mammary tumors with metastases in both the lung and liver. Transplantation of the tumors revealed metastases to lymph nodes that was enriched through serial transplantation. Through whole genome sequencing and RNAseq analysis we identified, and then confirmed *in vivo*, that Cyclin D1 was dysregulated in E2F5 conditional knockout mammary glands and tumors. Based on these findings, we propose that loss of E2F5 leads altered regulation of Cyclin D1, which facilitates the development of mammary tumors.

## Introduction

The mammary gland is a complex organ that undergoes dynamic changes during different stages of development. Analysis of transcriptional profiles at each developmental stage, including pregnancy, lactation and involution, have revealed unique gene expression changes^1,2^. One of the family of transcription factors that regulate these intricate transcriptional changes are the E2F transcription factors. The E2F transcription factor family consist of 9 members that can be divided into two groups based on their roles as a transcriptional activator or repressor^3-6^. E2Fs are best known for their role in cell cycle progression^7-10^. However, they are a functionally diverse group of transcription factors with numerous studies highlighting their role in apoptosis^11^, cell differentiation^12,13^, metabolism^14^ and development^15-19^. The role of E2Fs in development have been established through the characterization of single and compound E2F knockout mice^15-19^. Moreover, the role of E2Fs in mammary gland development were first characterized in E2F1, E2F2, E2F3 and E2F4 knockout mice^20^. Loss of E2F1, E2F3 and E2F4 resulted in mammary outgrowth delay and branching defects. However, these changes were not observed in in E2F2KO mice. In addition, E2F3 heterozygous mice demonstrated slight delay in involution. Given the high functional redundancy observed among E2Fs^21^, the extent of compensation between the activator E2Fs in the mammary gland developmental processes was explored. Using double knockout mice, this revealed that E2F2 can partially compensate for the loss of E2F1 but not E2F3 in mammary outgrowth in the double knockouts^22^. Although studies have characterized the roles of activator E2F1-3 and repressor E2F4 in mammary development and ductal morphogenesis, little is known whether the other repressor, E2F5, has a role in mammary development. Similar to E2F4, E2F5 is considered to be a transcriptional repressor and canonically functions to repress cell cycle progression^23,24^. Moreover, E2F4 and E2F5 share the most structural similarities among all the E2F members and have demonstrated functional redundancy^23^.

In addition to regulation of development and the well-known regulation of cell cycle, the E2F family has clear roles in cancer, including breast cancer. For instance, specific activator E2Fs have unique roles in mediating metastasis, roles that are dependent upon the context of the activating oncogene with noted differences in tumors driven by Myc, Neu or PyMT^25-29^. E2F functions extend beyond mediating cell cycle or transcriptional programs regulating metastasis with noted effects on vasculature, immune regulation and microenvironment^30^. In addition to transcriptional control of cell signaling pathways, E2Fs have the potential to regulate oncogenes, with well characterized effects both up and downstream of Myc ^6,31-33^ but with other oncogenes such as cyclin D1^34^ having been identified as potential targets through ChIP-Seq experiments^9,35,36^. Given the broad range of transcriptional targets regulated by the E2Fs^37^, it is not surprising that in the complex context of signaling in the tumor and tumor microenvironment that key targets are subject to E2F regulation.

The role of the E2Fs in development and function of the mammary gland, and the alterations to E2Fs altering tumor biology, have been described for the activator E2Fs but the repressor E2F5 has been understudied. This is partially due to the early lethality associated with the knockout of E2F5 resulting in hydrocephaly in prepubertal mice^19^, making phenotypic examination and experimental manipulation more difficult. Interestingly, the hydrocephaly is distinct from the E2F4 knockout phenotype^18^ but both combine to contribute to cell cycle^23^. The literature for E2F5 in cancer biology reveals a lack of consistency with some studies suggesting an inhibition of transformation^38^ while others note an oncogenic role^39-41^. Given the wide variety of target genes regulated by the E2Fs, it is likely that the role of E2F5 is largely tissue, cell type and context dependent, with roles that widely differ in tumor types. Based on gene expression data we have hypothesized that E2F5 plays an essential role in mammary gland development. To test this prediction, we have generated mice with a conditional ablation of E2F5 in the mammary gland. In these mice we have observed alterations to development and after a long latency, formation of metastatic tumors. Importantly, the transplantable tumors from these mice develop lymph node metastases, a trait not widely reported for other genetically engineered mouse models of breast cancer.

## Results

To begin to investigate a potential role for E2F5 in mammary development and function, we used a scRNAseq dataset^42^ that was clustered into the various stages, including nulliparious, lactating and involuting and tested for E2F5 expression. This revealed higher expression of E2F5 in lactating cells (Figure 1A). Interestingly, a comparison of developmental stages revealed that E2F5 was expressed in progenitor cells as well as the differentiated basal and luminal populations, a sharp contrast to the E2F1, a typical activator E2F (Supplemental Figure 1A). Given the expression in different cell types we assessed E2F5 expression in a survey of various cell types within the mammary gland^43^, using an involuting gland to have good representation of immune cells during remodeling. This revealed that E2F5 expression was present in the secretory alveoli as well as endothelial cells and fibroblasts (Figure 1B). While expression of E2F5 leads to plausible hypotheses about function, expression is not linked to activity. To assess this, we generated a signature for E2F5 by overexpressing E2F5 using an adenoviral vector in Human Mammary Epithelial Cells (HMECs), with increasing multiplicity of infection tied to E2F5 levels (Figure 1C). After infection, RNA was isolated to generate a gene expression signature using microarrays. HMECs infected with the E2F5 construct were compared to controls with a GFP construct and a signature was generated using a Bayesian approach. The up and down regulated genes in the signature (Figure 1D) were then tested for their predictive activity on a series of breast cancer cell lines where predicted activity was compare to protein levels from a Western blot, revealing that the signature was indeed predictive (Figure 1E). A mammary gland developmental dataset^1^was then limited to the E2F5 signature genes and clustered, revealing that E2F5 regulated genes stratified mammary developmental stages (Figure 1F). In addition, we tested genes with a fold change in a terminal end bud / duct dataset^44^ and found that nearly 10% of terminal end bud genes overlapped with E2F5 regulated genes (Supplemental Figure 1B). Thus, we hypothesized that E2F5 played several roles in the mouse mammary gland, including developmental and functional roles.

**Figure 1.**
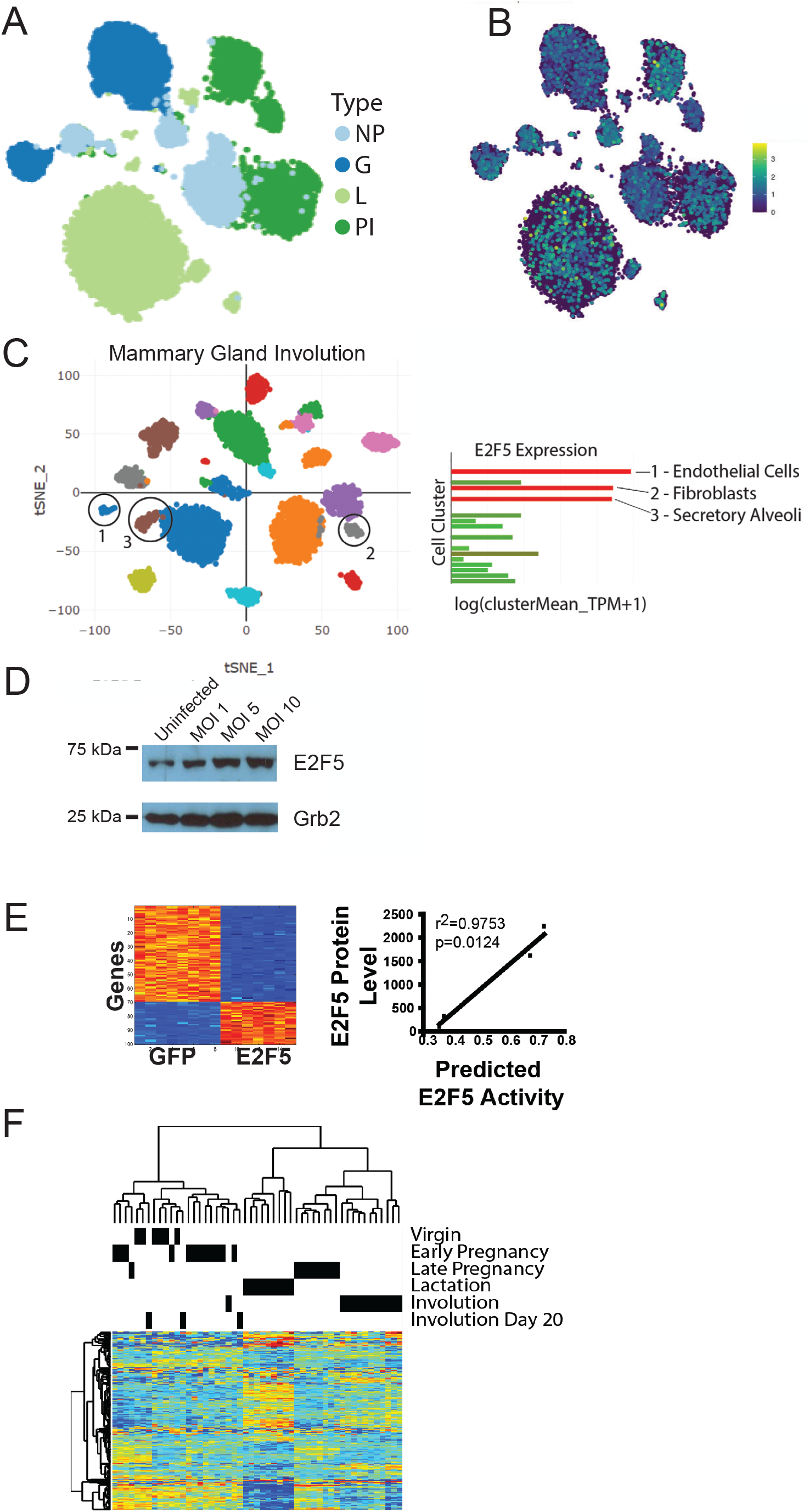
Prediction of a developmental role for E2F5 in the mouse mammary gland. Using a mouse mammary scRNAseq dataset that was split to various functional stages (A) including nulliparous (NP) gestation day 14.5 (G), lactation day 6 (L) and involution day 11 (PI). For each stage, the level of E2F5 expression was plotted (B). Using a separate scRNAseq dataset that was not sorted for epithelial cells, expression of E2F5 was examined across cell populations in the involuting mammary gland, revealing expression in endothelial cells, fibroblasts and alveoli (C). To generate a signature for E2F5 activity, HMECs were infected with increasing multiplicity of infection (MOI) for an adenovirus expressing GFP or E2F5. A western blot demonstrated increasing levels of E2F5 with increasing MOI (D). Generation of a signature for E2F5 activation revealed genes up (orange/red) and down (blue) regulated (E). The activity of the signature was predicted in several human breast cancer cell lines and was plotted against the observed levels in a western blot for E2F5 revealing correlation between the two. A mammary gland developmental dataset was limited to the E2F5 signature genes and was clustered, revealing that genes regulated by E2F5 stratified mammary developmental stages (F).

To directly test the role of E2F5 in mammary development we sought to use a knockout mouse model. With the lethality observed in the global knockout^19^, we generated a strain of mice with loxP sites flanking exons 2 and 3 of E2F5 (Figure 2A). Through breeding to the MMTV-Cre strain^45^ with expression limited to mammary epithelium^46^, this will result in a mammary specific knockout of E2F5 (Figure 2B). Prior to interbreeding with MMTV-Cre transgenics, the E2F5^flox/flox^ mice were backcrossed into the FVB background for 12 generations. As expected, introduction of MMTV-Cre to the E2F5^flox/flox^ strain resulted in excision, which was not complete since whole mammary glands were tested and Cre expression is limited to mammary epithelium (Figure 2C). Testing for a developmental role, we examined mammary gland outgrowth at 4 weeks of age where the littermate controls had terminal end buds that had driven outgrowth past the lymph node in the fat pad (Figure 2D). The E2F conditional knockouts (E2F5 CKO) however had development that lagged behind the controls and had only begun to reach the lymph node (Figure 2E). Quantification of these results revealed a marked delay in ductal extension (Figure 2F, p=0.0019). Other stages of mammary gland function were also assayed, with lactation occurring normally as assessed by histology and pup weight (data not shown). Involution had a slight delay that was apparent by the fourth day (Figure 2G-H). Latter stages of virgin development and involution revealed no significant delays. As the mice aged, we noted occasional regions of additional alveolar development. To test these effects we allowed a cohort of controls and E2F5 CKO mice to age to 12 months and examined wholemounts and histology. This revealed standard development in the controls and surprising overgrowth resembling lactation in the virgin E2F5 CKO mice (Figure 2I-J).

**Figure 2.**
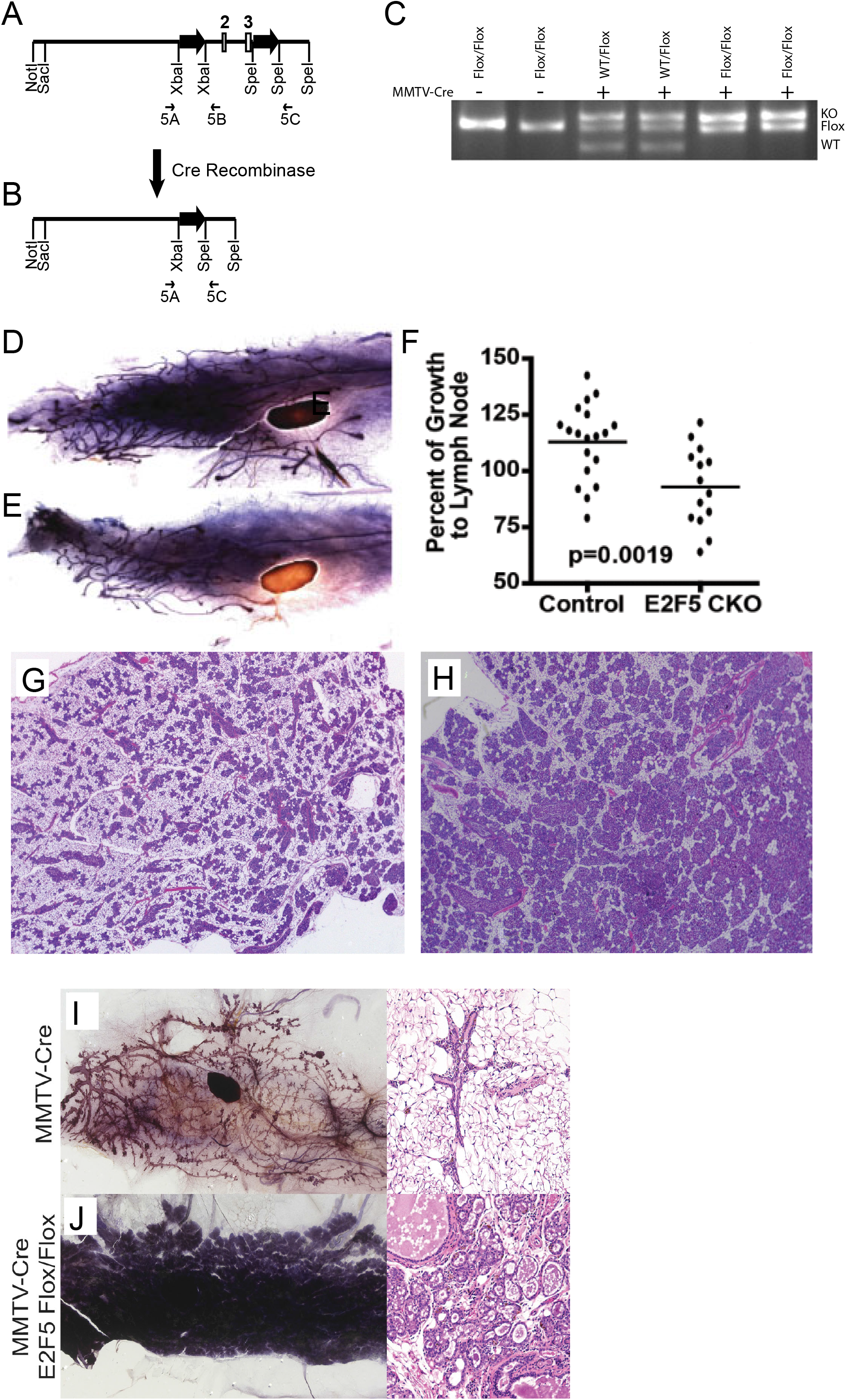
Developmental delays with conditional loss of E2F5. A gene targeting strategy to flank exons 2 and 3 of E2F5 with loxP sites was employed with genotyping primers shown (A). With the introduction of Cre Recombinase, exons 2 and 3 are lost (B), resulting in a tissue specific knockout. Using the indicated primers, the wild type (WT), loxP flanked (flox) and knockout (KO) alleles were detected with and without expression of MMTV-Cre in the mammary glands of 8 week old virgin mice (C). Examining ductal extension through wholemounts at 4 weeks we examined controls (E2F5^flox/flox^) (D) and E2F5 CKO mice (E), revealing a delay in outgrowth. This delay was quantified revealing a significant outgrowth delay, 0=0.0019 (F). A delay in involution was also noted with the 4^th^ day after pups were removed from the dams having the most visible delay. Relative to E2F5^flox/flox^ controls (G), the E2F5 CKO mice had more secretory alveoli (H). After mice aged to 12 months, virgin mammary glands were assessed by both wholemount and histology. Relative to the MMTV-Cre controls (I) with their somewhat spiked ductal appearance, the E2F5 CKO mammary glands resembled a lactating mammary gland with alveoli engulfing the entire fat pad (J).

With the overgrowth of the virgin mammary gland we carefully observed the E2F5 CKO mice for mammary tumor development. After a long latency, virgin E2F5 CKO mice developed focal mammary tumors while the MMTV-Cre control line did not (Figure 3A). Multiparous E2F5 CKO mice were noted to have a slight reduction in tumor latency, potentially due to more widespread Cre expression or E2F5 effects during pregnancy, lactation or involution. The tumors that formed exhibited varied histological patterns, with a broad spectrum of subtypes reminiscent of those noted in other heterogenous strains such as MMTV-PyMT^27,47^(Figure 3B-E). These tumors were highly metastatic with 74.2% of tumor bearing mice developing an average of 2.31 pulmonary metastases. In addition, 20% of these mice also developed liver metastases (Figure 3F). An example of the pulmonary metastases is shown with several regions of the lungs magnified, demonstrating clear differences in the pathology of the metastatic lesions (Figure 3G). In addition to pulmonary and liver metastases, other occasional metastatic lesions were noted in other locations, including lesions on the thoracic wall, lymph nodes and in the intraperitoneal cavity. With loss of E2F5 resulting in tumor formation in the mouse model, we hypothesized that E2F5 function may be related to human breast cancer development or outcomes. Testing the E2F5 signature in human breast cancer allowed us to stratify patients to high / low quartiles of E2F5 activity. Examining survival of these patients revealed that low E2F5 activity was associated with a reduction in survival rates (Figure 3H). Moreover, at the single gene level, low levels of E2F5 expression in subtypes of breast cancer was linked to poor outcomes (Supplemental Figure XXX).

**Figure 3.**
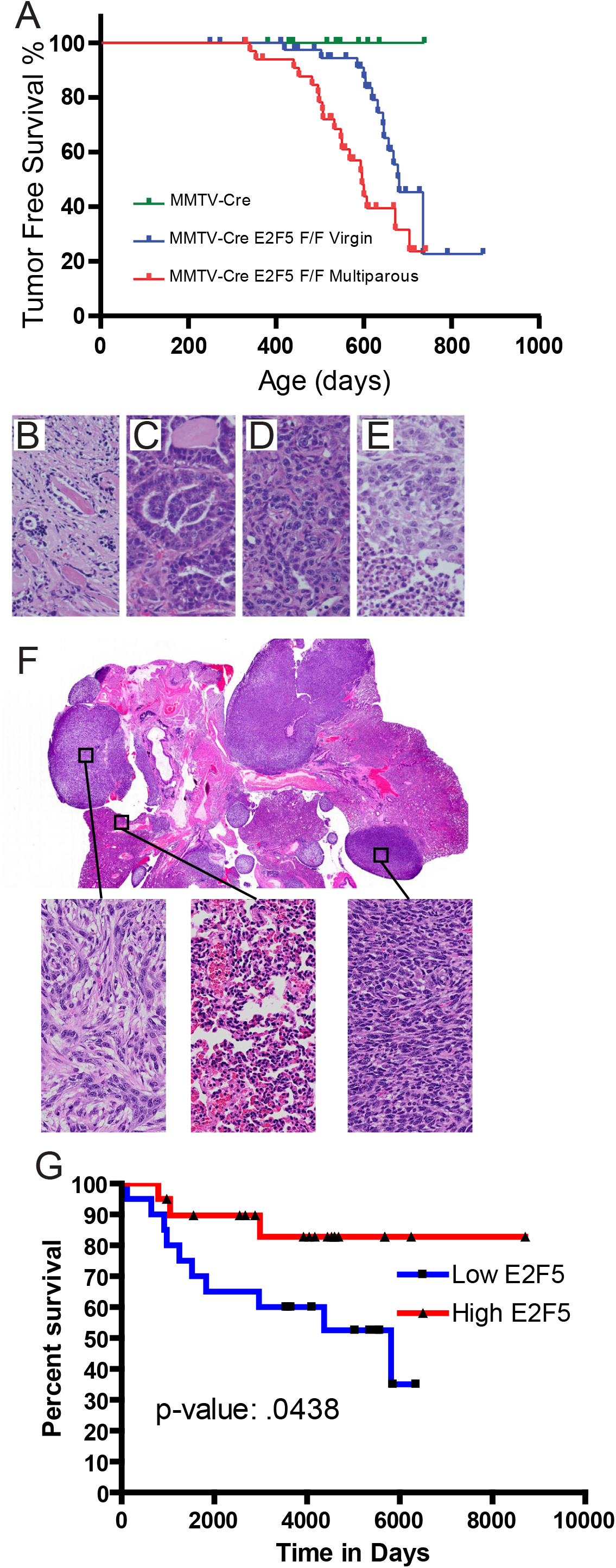
Metastatic tumor formation in mice lacking E2F5 in the mammary epithelium. Regular palpation of the mammary glands revealed the onset of tumor formation in E2F5 CKO virgin and multiparous mice, but not in the control MMTV-Cre line (A). The resulting tumors were histologically diverse with numerous patterns noted (B-E). The tumors were also metastatic, a pulmonary section reveals numerous metastatic lesions at both low and high power (F). Applying the E2F5 signature to human breast cancer stratified patients to high/low quartiles. These results were consistent with the mouse model as low E2F5 activity was associated with worse survival.

The development of metastases that reflect human breast cancer is an attractive component of this new mouse model of breast cancer but that is offset by the extended latency. During necropsy of the tumor bearing mice, small portions of the tumor were viable frozen. These small tumor fragments were then thawed and directly implanted into mammary glands without passing through tissue culture. MMTV-Cre mice were used as transplant recipients to prevent immune effects associated with the MMTV promoter enhancer^48^. 18 frozen and one fresh tumor were implanted into a small pocket created in the abdominal mammary gland of recipient mice, rapidly resulting in overt tumor formation. Strikingly, it was repeatedly noted that in addition to the primary tumor these mice developed a secondary metastatic tumor in the axial lymph node in 12 of the 19 lines at low penetrance (Figure 4A). Examining the histology of these metastatic tumors revealed the presence of both lymphatic and tumor tissue (Figure 4B). Pan-cytokeratin staining for epithelial derived tumor cells revealed nests of tumor cells within the axial lymph node (Figure 4C). Hypothesizing that the metastasis was occurring through the lymphatic vasculature, we sectioned across the region between the primary tumor and axial lymph node from Figure 4A and stained for podoplanin, a marker of lymphatic vessel walls. This revealed staining of the lymphatic vasculature with tumor cells lodged within (Figure 4D). Control transplants of MMTV-Neu and MMTV-PyMT tumors did not result in lymph node metastasis (data not shown). Given prior enrichment strategies for breast metastases^49-51^, we then implanted the lymph node metastases to the mammary gland of new recipient mice, resulting in an enrichment of lymphatic metastases (Figure 4E).

**Figure 4.**
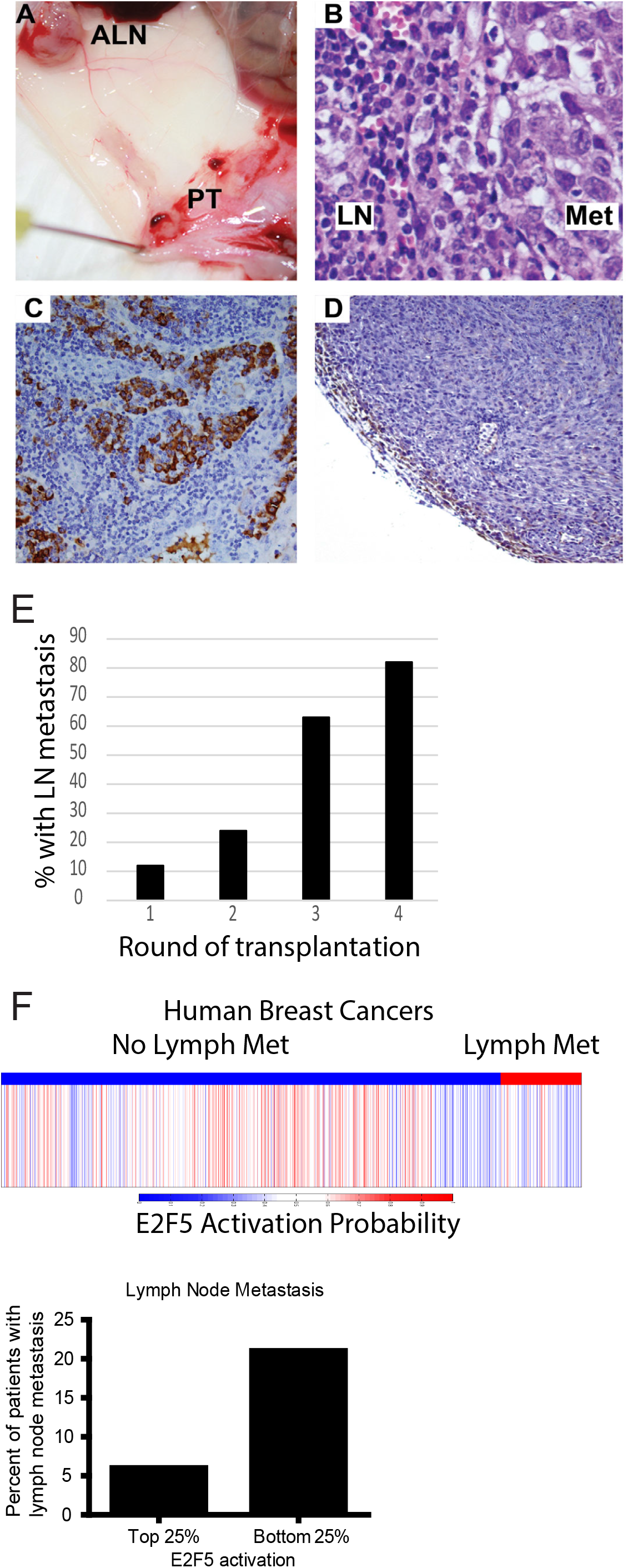
E2F5 CKO tumors develop lymphatic metastasis. Implantation of E2F5 CKO tumors into FVB MMTV-Cre recipients resulted in tumor formation. The primary tumor (PT) has been excised but a tumor has also formed in the region of the axial lymph node (ALN) (A). Cross section of the lymph node reveals both lymph (LN) tissue and metastatic (MET) cells (B). Staining with a pan-cytokeratin antibody reveals nests of metastatic cells throughout the lymph node (C). Cross section and staining of the lymphatic vessel from (A) for podoplanin reveals a lymphatic vessel containing tumor cells (D). Serial transplantation of various E2F5 CKO lines from lymph node to mammary gland resulted in the enrichment of the lymphatic metastasis phenotype (E). Examining human breast cancer with known lymph node status for the E2F5 activation gene signature revealed that metastatic tumors had lower E2F5 activity. Splitting the samples into quartiles, the samples with the lowest levels of E2F5 were most likely to have lymph node metastasis (F).

The diagnosis of human breast cancer uses TMN (tumor, metastasis, node status) staging, with lymph node status as one of the key elements. After observing lymphatic metastasis in our mouse model, we sought to test for a role for E2F5 in human lymphatic metastasis. To address this, we predicted E2F5 activity using our signature in human breast cancer with known nodal status. Stratification of breast cancers into +/-lymph node metastasis revealed that those tumors with lymph node metastases had lower levels of E2F5 (Figure 4F). Examining the upper and lower quartiles for E2F5 activity revealed almost a four-fold increase in lymphatic metastasis in the lower quartile of E2F5 activity.

To explore the mechanisms by which E2F5 may regulate tumor development and progression we used an integrated approach analyzing both transcriptomic and whole genome sequence data. Bulk tumors were used for RNAseq and the results were clustered together with three commonly used mouse models of breast cancer to examine similarities and differences of the models. This analysis revealed that E2F5 clustered with subsets of these tumors with the heterogeneity that was observed histologically being reflected in the gene expression patterns (Figure 5A). Geneset enrichment analysis (GSEA) of these tumors revealed a similarity to the luminal subtype of human breast cancer (Figure 5B-C).

**Figure 5.**
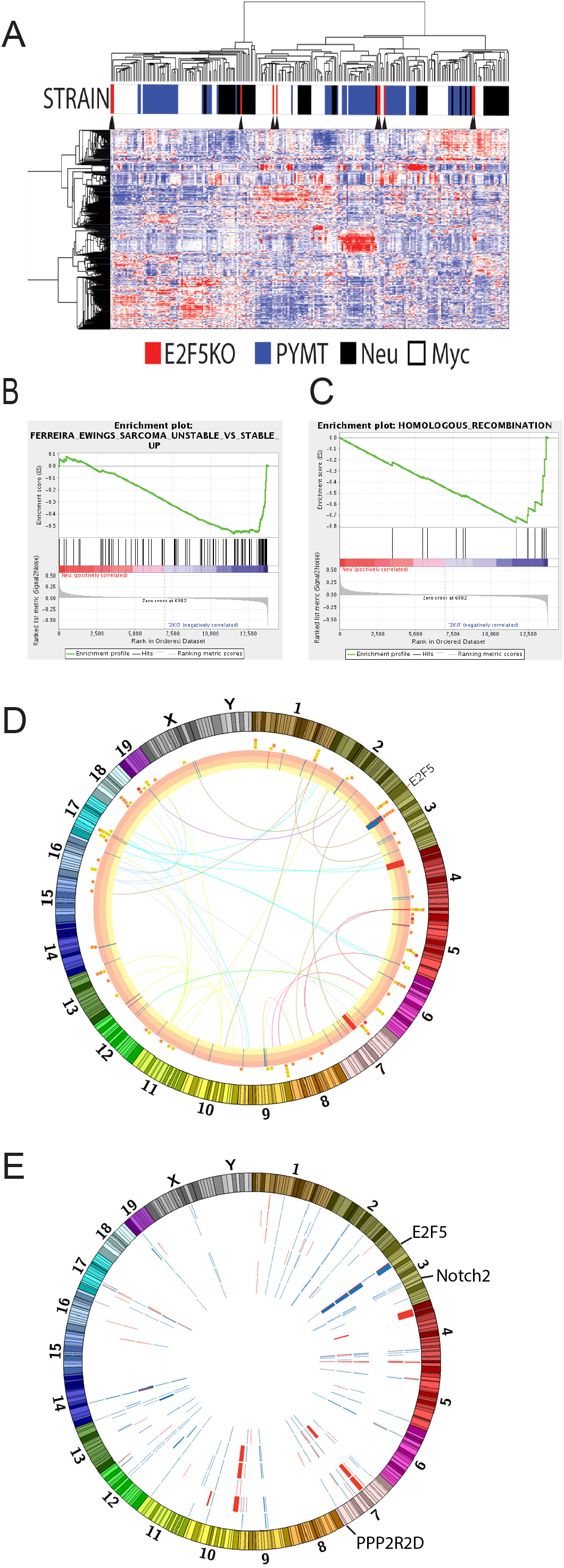
Transcriptomic and sequence analysis of the E2F5 CKO tumors. Examining RNAseq for E2F5 CKO tumors relative to the MMTV-PyMT, Neu and Myc strains revealed that the E2F5 CKO tumors were just as transcriptionally diverse and clustered into several distinct groups (A). Examination of GSEA pathways revealed several distinctions between the E2F5 and other tumor models (B-C). Whole genome sequencing of E2F5 CKO tumors revealed SNV, CNV and translocations for the example tumor shown (D). Comparison of CNV events from 6 E2F5 CKO tumors revealed a surprising number of shared events, including alterations to Notch2 and PPP2R2D (E).

In addition to gene expression, we examined whole genome sequence for 6 tumors with diverse histopathology. TCGA best practices were followed for analysis. A representative circos plot for one of the tumors is shown with chromosomes around the periphery. Moving towards the interior, single nucleotide variants (SNV) are shown as points with color reflecting putative impact, followed by copy number variants (CNV) with deletions and amplifications shown with color blocks and then in the center the translocations and inversions are shown with lines (Figure 5D). To examine shared CNV events, an additional circos plot was generated with each of the 6 tumors represented by a ring inside of the chromosomes (Figure 5E). This analysis revealed a surprising number of common events for both deletion and amplifications. This includes CNV for key genes such as Notch2, ODR4, ITLN1 and SDHD. SDHD is part of the succinate dehydrogenase enzyme complex and is associated with a range of familial cancers^52^ and a search of the 2015 TCGA breast cancer database in cBioportal reveals 7% of tumors with SDHD alterations.

To better understand transcriptional profiles of E2F5 CKO tumors, RNAseq was performed on MMTV-Cre mammary glands, E2F5 CKO mammary glands, E2F5 CKO mammary tumors and tumor cell lines derived from the E2F5 CKO tumors. Differential gene expression analysis was performed between MMTV-Cre and E2F5 CKO mammary glands and MMTV-Cre mammary glands and E2F5 CKO tumors/cell lines. To identify gene expression changes driven by E2F5 loss, we composed a list of genes that are differentially regulated in both E2F5 CKO mammary glands and tumors relative to MMTV-Cre mammary glands. These genes were filtered using several criteria including fold change, percent alteration in human breast cancer and known E2F target status from public ChIPseq data. Based on these factors, 4 candidates (Rad51, Sphk1, Kif20a and Cyclin D1) were chosen for validation with qRT-PCR. In line with the differential gene expression analysis, Cyclin D1 (Supplemental Figure X) and the other three genes (data not shown) demonstrated increased expression in E2F5 CKO mammary glands relative to MMTV-Cre mammary glands through qRT-PCR. Given that our E2F5 signature was derived from overexpression of E2F5, we hypothesized that this should result in a decrease in cyclin D1 levels, with this finding confirmed in gene expression data (Supplemental Figure X). However, the most striking difference was seen in Cyclin D1 where there was a significant increase in E2F5 CKO tumors relative to control in the RNAseq data (Figure 6A), confirmed with a 15 fold increase in *cyclin D1* levels observed through qRT-PCR (Figure 6B). Given prior reports of altered levels of Cyclin D1, D2 and D3 in mouse mammary tumor models^53^, we examined this family of genes in Wnt-1, Neu and the E2F5 CKO tumors. This revealed that levels of Cyclin D1 were consistent in all three models but both E2F5 CKO and Neu induced tumors had only only Cyclin D1 elevated in the majority of tumors (Figure 6 C).

**Figure 6.**
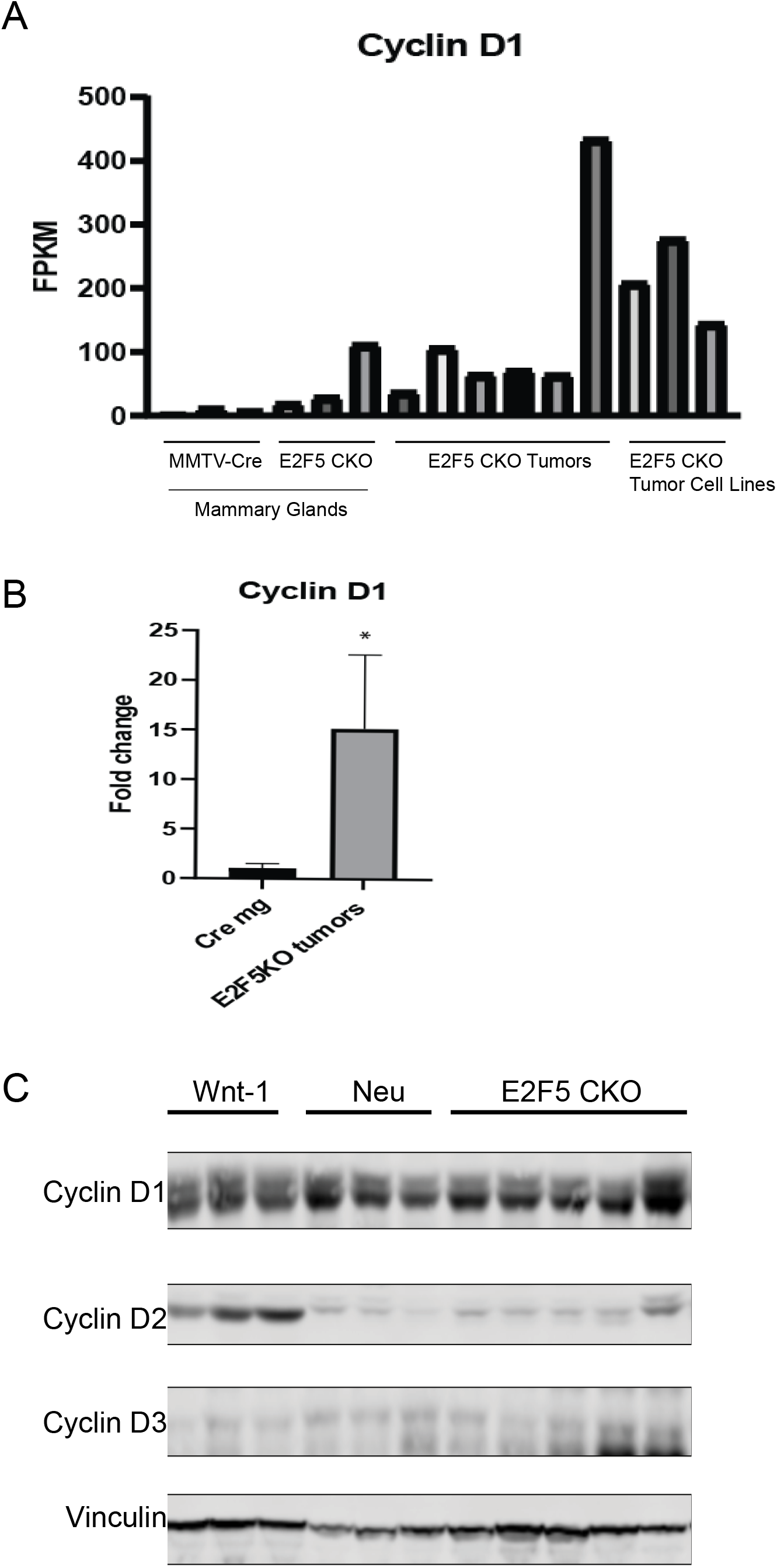
Cyclin D1 is activated in E2F5 CKO tumors. Using a filtering strategy to determine genes involved with E2F5 CKO tumor formation, we identified Cyclin D1. Comparison of mammary gland expression of cyclin D1 through RNAseq revealed an upregulation in mammary glands, tumors and more extensively in the cell lines derived from the tumors (A). Validation of these results through qRT-PCR for cyclin D1 in MMTV-Cre and E2F5 CKO mammary glands revealed a 15 fold upregulation of cyclin D1 in the pre-tumor mammary glands. Examination of Wnt1, Neu and E2F5 CKO tumors revealed that each strain had elevated Cyclin D1 but only Wnt1 tumors had upregulation of Cyclin D2 (C).

## Discussion

Here we have used gene expression data to predict a role for the E2F5 transcription factor, commonly thought of as a traditional repressor, in both mammary gland development and tumor formation. Testing the role of E2F5 required generation of a conditional deletion due to the early lethality of the whole body knockout due to developmental abnormalities^19^. Here we created a conditional deletion of E2F5 through generation of a floxed E2F5 allele interbred with MMTV-Cre mice, which we previously characterized to have mammary epithelial specific expression of Cre recombinase^45,46^. Surprisingly only minor mammary gland defects were noted with the mammary specific loss of E2F5 but, after an extensive latency, metastatic mammary tumors developed. Notably, these tumors metastasized to the lymph node and this property was enhanced with serial transplantation. Mechanistically, we demonstrate an elevated level of Cyclin D1 with similarities to the MMTV-Neu tumor model.

During generation of the E2F5 gene signature we overexpressed E2F5 in HMECs and collected RNA 18 hours after infection, a standard procedure that allows immediate transcriptional events to be assayed^54,55^. The conventional role for E2F5 is thought to be a transcriptional repressor^7,23^, although several more recent publications have demonstrated a role as an activator in diverse settings include zebrafish^56^ and cervical cancer^57^. Our results demonstrate that E2F5 can function both as an activator and as a repressor in mammary epithelial cells, dependent upon the gene. This is consistent with prior work suggesting that the context of E2F binding is an essential component in determining function^58^.

While the predictive bioinformatic data strongly suggested an essential role for E2F5 in several stages of mammary development, from regulation of genes expressed in the terminal end bud to stratification of the major stages of mammary development by E2F5 signature genes, the observed phenotypes were only minor delays in outgrowth and involution. These were transient delays with mice retaining full function of the mammary gland in allowing the dam to raise healthy pups. Prior work has shown a range of individualized roles for E2F1-4 in mammary development with E2F4 having the most severe developmental effects and a near inability to rear pups^20^. However, consistent with the notion that the E2Fs can compensate for each other^21^, we recently functionally demonstrated that E2F compensation occurs during mammary gland development^22^. This is reinforced by the more severe effects seen in E2F4/5 double knockouts relative to the individual knockouts in cell cycle progression, moreover the double knockouts suffered from neonatal lethality while the individual knockouts were viable at birth^23^. Together our data and the literature suggest that E2F5 functions to regulate development in a context dependent manner with other E2Fs compensating for loss of any one E2F.

Although developmental alterations were minimal, E2F5 CKO mice spontaneously developed mammary tumors. Similar to human breast cancer, mammary tumors arising in E2F5 CKO mice demonstrated diverse morphology. Furthermore, the prolonged tumor latency in the E2F5 CKO model is similar to human breast cancers, where the majority of cancers occur in older postmenopausal women. Given that majority of the transgenic mouse models develop tumors with at a shorter latency^59^, E2F5 CKO mice may be a useful for model for studying age-related changes in breast cancer. The prolonged tumor latency also suggests that, in addition to the loss of E2F5, other genetic events need to accumulate prior to tumor initiation. Remarkably, many of these CNV events were conserved across the model system, despite tumors being histologically diverse.

In the study of metastasis in genetically engineered mouse models, pulmonary metastasis is commonly noted and we found E2F5 CKO mice developed pulmonary metastasis. Interestingly, we discovered that E2F5 CKO mammary tumors transplanted into the abdominal mammary fat pad had a propensity to metastasize to the axillary lymph node. Given that axillary lymph nodes are most commonly the first site of metastasis in human breast cancer, we sought to enrich the ability of E2F5 CKO tumors to metastasize to the axillary lymph node. Using a serial transplantation technique to re-transplant the axillary tumor into the abdominal mammary fat pad, we generated a syngeneic transplantation model that develop lymph node metastasis within one month of transplant and with >80% penetrance. Current mouse models of breast cancer rarely metastasize to the lymph node. Therefore, this model of enriched lymph node tumors is unique and can provide a unique resource to examine the mechanisms driving lymph node metastasis.

Integrating our bioinformatic analysis and *in vitro* studies, we identified Cyclin D1 as a potential mediator of tumor development and progression in E2F5 CKO tumors. Cyclin D1 is commonly amplified and/or overexpressed in human breast cancer^60^. Moreover, overexpression of Cyclin D1 in the mouse mammary gland results in mammary tumor development, supporting a key role in tumorigenesis^34^. Based on gene expression analysis, E2F5 expression is inversely correlated with Cyclin D1, suggesting that E2F5 may be negatively regulating Cyclin D1. To investigate the role of Cyclin D1 in E2F5 CKO tumors, we examined Cyclin D1, D2 and D3 levels in E2F5 CKO tumors compared to MMTV-Wnt-1^61^ and MMTV-Neu^62^ tumors. Our analysis revealed similar levels of Cyclin D1 expression across all three models. However, Cyclin D2 and D3 levels were lower in the MMTV-Neu and MMTV-E2F5KO models compared to the MMTV-Wnt model. Previous work has shown that Cyclin D1 loss does not affect MMTV-Wnt tumor development because this strain also expresses Cyclin D2 and is able to compensate for Cyclin D1 loss^53^. In contrast, since MMTV-Neu mainly express Cyclin D1, the loss or inhibition of Cyclin D1 inhibited tumor development as demonstrated in several studies^53,63,64^. However, Zhang *et al*. demonstrate that loss of Cyclin D1 in MMTV-Neu tumors only delayed tumor latency since Cyclin D3 is able to compensate for Cyclin D1 loss^64^. Thus, there are contradicting results in whether tumor development still occurs in Cyclin D1 deficient MMTV-Neu mice as a result of Cyclin D3 compensation. However, these discrepancies do not undermine the fact that Cyclin D1 is critical for tumor development and under normal circumstances is the main member of D-type cyclins to initiate tumorigenesis in MMTV-Neu. Furthermore, Cyclin D1 expression in MMTV-Neu tumors is mediated by E2F1^63^. Other studies have also demonstrated that E2F1 and E2F4 can directly bind to and regulate Cyclin D1 expression^65^. Given the functional redundancy and shared binding motif between E2F family members, it is likely that E2F5 can also regulate Cyclin D1 expression. Taken together, we propose that loss of E2F5 in the mammary gland leads to deregulation of Cyclin D1, contributing to tumor development and progression. Given the wide range of SNV and CNV alterations, we suggest that this occurs in a complex mutational environment with numerous other pathways. Although there is evidence suggesting that E2F5 may be directly Cyclin D1 expression, it is also possible that disruption of E2F5 leads to dysregulation of other targets that can result in Cyclin D1 expression.

In this study, we have identified a functional novel role of E2F5 as a tumor suppressor. This is consistent with prior literature suggesting E2F5 has varied roles, including as a tumor suppressor^24^. E2F5 CKO mice develop histologically diverse mammary tumors and metastatic lesions after a prolonged latency. Given the lack of lymphatic metastasis genetically engineered mouse models, the E2F5 CKO syngeneic transplantation model can be significant resource to studying the mechanism of lymphatic metastasis. Finally, given the prolonged tumor latency and diverse morphology of E2F5CKO tumors, the E2F5 CKO mice can be a favorable model when studying age-related cancer changes.

## Material and Methods

### Animal generation

All animal husbandry and use was in compliance with local, national and institutional guidelines. Ethical approval for the study was approved by Michigan State University Animal Care & Use Committee (IACUC) under AUF 06/18-084-00. E2F5 CKO mice were generated by flanking exons 2 and 3 of the E2F5 gene with loxP sites. E2F5^flox/flox^ mice were interbred with MMTV-Cre mice^45^. Mice were monitored weekly for tumor development. The endpoint for primary tumor was 2000 mm^3^.

### Cell culture

Human mammary epithelial cells were cultured in Mammary Epithelial Cell Basal Medium (ATCC, Manassa, Virginia #PCS-600-030) supplemented with rH-insulin (5 ug/mL), L-glutamine (6 mM), epinephrine (1 μM), apo-transferrin (5 μg/ml), rH-TGF-α (5 ng/ml), extractP (0.4%) and hydrocortisone hemmisuccinate (100 ng/ml). BT549 were cultured in RPMI supplemented with 10% FBS (Gibco #10437028) and 1% Antibiotic-Antimycotic reagent (Thermo Fisher, Waltham, MA, USA #15240062). MDA-MB-231 were cultured in DMEM supplemented with 10% FBS (Gibco # 10437028) and 1% Antibiotic-Antimycotic reagent (Thermo Fisher, Waltham, MA, USA #15240062).

### E2F5-regulated genes

Human Mammary Epithelial cells were infected with adenovirus expressing E2F5 or GFP. Cells were collected eighteen hours after infection. Total RNA was extracted using Qiagen RNeasy Mini kit. RNA was used with Affymetrix Human Genome U133 chip to generate gene expression data. RMA algorithm was used to normalized microarray dataset. Significance Analysis of Microarray was applied to the dataset to identify differentially expressed genes in HMEC-E2F5.

### Pathway analysis

Single Sample Gene Set Enrichment Analysis and Gene Set Enrichment Analysis were performed on Broad Institute Genepattern interface. Normalized RMA data was used as input for microarray data and normalized TPM data was used as input for RNA-seq data. E2F5 activation signature was generated from genes that were upregulated and downregulated in HMEC overexpressing E2F5 using standard methods^54^. E2F5 target geneset and other genesets used were derived from mSigdb.

### Histology

For wholemount analysis, abdominal mammary fat pads were excised and placed on glass slides. The slides were incubated in acetone for 24 hours, rehydrated through an ethanol progression and stained in Harris’ Modified Hematoxylin for 24 hours. The slides were destained in acidified ethanol. Slides were dehydrated and mounted with permount. To evaluate mammary outgrowth, the distance from the nipple to the leading edge of the epithelium and the distance from the nipple to the midpoint of the thoracic lymph node were measured. Samples for histology were fixed in 10% formalin and submitted to Michigan State University Pathology lab.

### Microarray analysis

Significant Analysis Microarray was used for differential gene expression in microarray data. The following published microarray datasets were used for analysis: terminal end bud and duct (GSE2988) and mammary gland developmental stages (GSE12247).

### RNA-sequencing

Flash frozen tumor pieces were homogenized using Fisher Homogenizer 150 (Thermo Fisher, Waltham, MA, USA). Total RNA was isolated using QIAGEN RNeasy Midi Kit (Hilden, Germany #75142) with the manufacturer’s protocol. RNA concentration was measured by Qubit and Agilent 2100 Bioanalzyer. RNA samples with RIN >7 was used for library preparation using the Illumina Tru-Seq stranded total RNA kit. RNA library was sequenced to a depth of >20M reads/sample with paired end 150 base paired reads on Illumina NovaSeq 6000. Adaptors were removed from reads using Trimmomatic v0.33. Quality control was performed using FastQC v0.11.5. Reads were aligned and mapped using STAR [35]. RSEM was used to quantify and normalize reads [36]. Differential gene expression analysis was performed using EdgeR.

### Clustering

Unsupervised clustering was performed using Broad Institute’s Morpheus interface.

### Immunoblotting

To extract RNA from tissue, samples were homogenized using mortar and pestle in liquid nitrogen. Sample were lysed in TNE lysis buffer (0.05 M Tris HCl pH 8.0, 0.15 M NaCl, 2 mM EDTA, 0.01 N NaF and 1% NP40) with proteinase inhibitor (1 M Na3VO4, 58 μM PMSF, 10 μg/ml aprotinin and 10 μg/ml leupetin) for 1 hour on ice with constant agitation. Protein was quantitated using BCA (Thermo Scientific, Waltham, MA, USA #23225) and then boiled at 100°C for 5 min. Samples were loaded onto a 8-12% polyacrylamide gel. Separated protein was transferred onto a Immobilon-FL PVDF membrane (Millipore Sigma, Burlington, MA, USA #IPFL00010). Membranes were blocked in 5% milk in TBS with 0.1% Tween-20 (TBS-Tween) for 1 hour and then incubated in primary antibody overnight at 4°C. Following three washes in TBS-Tween the membrane was incubated in the appropriate antibody at a dilution 1:10,000 in 5% milk in TBS-Tween for 1 hour at room temperature. Membranes were washed 3x in TBS-Tween and imaged on LI-COR Odyssey imaging system (LI-COR Biosciences, Lincoln, NE, USA). The following antibodies were used: 1:1000 ERK (C-9), 1:100 E2F5 (C-8) from Santa Cruz Biotech (Santa Cruz, CA, USA), 1:1000 AKT, 1:1000 phospho-AKT, 1:4000 Vinculin from Cell Signaling Technology (Boston, MA, USA), 1:1000 phospho-ERK, 1:2000 Cyclin D1, 1:2000 Cyclin D3 from Thermo Fisher (Waltham, MA, USA), 1:1000 Cyclin D2 from Proteintech (Rosemont, IL, USA).

### Quantitative RT-PCR

Flash frozen tumor pieces were homogenized using Fisher Homogenizer 150 (Thermo Fisher, Waltham, MA, USA). Total RNA was isolated using QIAGEN RNeasy Midi Kit (#75142; Hilden, Germany) with the manufacturer’s protocol. Quantitative RT-PCR was performed using Luna Universal One-Step RT-qPCR kit (New England Biolabs, Ipswich, MA, USA) according to manufacturer’s protocol using Agilent Mx3000P instrument. Primers were designed using Primer Bank tool (https://pga.mgh.harvard.edu/primerbank/). The following primers were used (5’ to 3’): Rad51 forward, *TGTTGCTTATGCACCGAAGAA*; Rad51 reverse, *GCTGCCTCAGTCAGAATTTTGT*; KIF20A forward, *CAGCGGGCTTACTCTCTGATG*; KIF20A reverse, *GTCTGACAACAGGTCCTTTCG*; Sphk1 forward, *ACTGATACTCACCGAACGGAA*; Sphk1 reverse, *CCATCACCGGACATGACTGC*; CCND1 forward, *TGACTGCCGAGAAGTTGTGC*; CCND1 reverse, *CTCATCCGCCTCTGGCATT*; Gapdh forward, *AGGTCGGTGTGAACGGATTTG*; Gapdh reverse, *TGTAGACCATGTAGTTGAGGTCA*. Primer efficiency was 90-110% for all primers used. Delta-delta CT method was used for fold change analysis.

### Mammary fat pad transplantation

E2F5 CKO mammary tumors were harvested and stored in DMEM with 20% FBS and 10% DMSO at −80°C before long term storage in liquid nitrogen vapor phase. Tumors were thawed and orthotopically implanted into the abdominal mammary gland of 6-to-10-week old MMTV Cre female mice. Mice were palpated 2x a week for mammary tumor formation. When the tumor size reached 2000 mm^3^, samples were harvested for further analysis.

### <u>*Whole genome sequence*</u> analysis

WGS sample reads were concatenated and quality checked using FastQC. Sample reads had adapter ends trimmed off using Trimmomatic and again checked for quality using FastQC. Paired WGS reads were aligned to the GRCm38-mm10 reference genome using Burrows-Wheeler Aligner (BWA/0.7.17). Read groups were added using Picard (picard/2.22.1). Reads were then indexed and sorted using SAMtools (SAMtools/1.11). Duplicate reads were removed using Picard. Final .bam files had SNVs called using the consensus of Mutect2, SomaticSniper, and VarScan. Translocations, inversions, and CNVs were called using the consensus of Delly and Lumpy for each tumor sample. Annotations of all samples was done using SnpEff.

### Statistical analysis

All statistical comparisons are performed with an unpaired students two-tailed, unpaired t-test.

